# **FusionCatcher** – a tool for finding somatic fusion genes in paired-end RNA-sequencing data

**DOI:** 10.1101/011650

**Authors:** Daniel Nicorici, Mihaela Şatalan, Henrik Edgren, Sara Kangaspeska, Astrid Murumägi, Olli Kallioniemi, Sami Virtanen, Olavi Kilkku

## Abstract

`FusionCatcher` is a software tool for finding somatic fusion genes in paired-end RNA-sequencing data from human or other vertebrates. `FusionCatcher` achieves competitive detection rates and real-time PCR validation rates in RNA-sequencing data from tumor cells. Fusion-Catcher is available at http://code.google.com/p/fusioncatcher/.

## 1 Introduction

Fusion genes are regarded as one of the common driver events behind tumor initiation and progression. Several fusion genes have also been found to be recurrent in multiple cancers. For example DNAJB1-PRKACA fusion was found, using `FusionCatcher`, in 100% of fibrolamellar hepatocellular carcinoma patients [15]. Also, fusion genes are good targets for therapeutic applications and personalized medicine [14].

One of the challenges in finding fusion genes in RNA-sequencing dataset is the very high rate of false positives which makes challenging their validation in the wet-lab. For example ChimeraScan[20] finds 28,399 fusion genes in four cancer breast cell lines[18] whilst:

- Cancer Genome Project lists 7,150 known fusion genes^1^[22, 21], and
- Mitelman Database of Chromosome Aberrations and Gene Fusions in Cancer lists 2,094 known fusion genes^2^ [24].

in thousands of cancer cells/patients.

Throughout the paper we refer to somatic fusion genes, or shorter fusion genes, as the fusion genes which are found in majority of times in diseased cells/samples/patients. Therefore the expectation here is that number of fusion genes found in healthy samples is zero or very close to zero.

Thus the main goals of `FusionCatcher` are: (i) good real-time PCR validation rate (i.e. precision) which makes practical the validation of candidate fusion genes, and (ii) good detection rate (i.e. sensitivity) of fusion genes.

Here, we present `FusionCatcher`, which is a software tool for finding novel and known somatic fusion genes in paired-end RNA-sequencing data in diseased samples from vertebrates which have annotation data available in Ensembl database[10].

## 2 Methods

First, on the RNA-sequencing data is performed some pre-processing and filtering. Quality filtering of reads is performed, by:

- removing reads which align on ribosomal/transfer RNA, mitochondrial DNA, HLA genes, or known viruses/phages/bacteria genomes,
- trimming the reads which contain adapters and poly-A/C/G/T tails,
- clipping the reads based on quality scores, and
- removing the reads which are marked as bad quality by Illumina sequencer.

In an RNA-sequencing experiment, the RNA (which is converted to complementary DNA) is sequenced usually using next-generation sequencing platforms. Therefore, `FusionCatcher` is performing most of the data analysis at the RNA level (i.e. transcriptome) by aligning the sequencing reads, as single reads, on transcriptome using Ensembl genome annotation[10] and Bowtie aligner[8]. Furthermore, the reads are mapped on genome, using the Bowtie aligner, for filtering purposes and the reads which have a better alignment (i.e. fewer mismatches) at the genome level will have their transcriptome mappings removed. Similarly, the reads, which map simultaneously on several transcripts of different genes, have their mappings on transcriptome removed, and for the corresponding genes a sequence similarity score is computed using these read counts. The unmapped reads, which are the reads which passed the quality filtering and do not map on the transcriptome and the genome, are kept for further analyses.

The reads mapping on the transcriptome are used further to build a preliminary list of candidate fusion genes by searching for pairs of genes, such that for each pair of genes (gene A, gene B) one read maps on gene A’s transcripts and its paired-read maps on gene B’s transcripts. From the preliminary list of candidate fusion genes are removed the pairs of genes, using known and novel criteria, which make biological sense, such as:

- both genes are known to be the other’s paralog in Ensembl database,
- a gene is known to be the other’s pseudogene in Ensembl database,
- a gene is known micro/transfer/Y/7SK/small-nuclear/small-nucleolar RNA,
- fusion is known *a priori* to be a false positive event (e.g. ConjoinG database [12], conjoined HLA genes [13],
- it has been found previously in samples from healthy persons, like for example from Illumina Body Map 2.0 RNA-sequencing data[23] and an in-house RNA-sequencing database of healthy samples,
- both genes are overlapping each other on the same strand according to one of the public known databases, such as Ensembl, UCSC, or RefSeq databases, and
- pair of genes which have a very high count of reads mapping simultaneously on both of genes forming a fusion.

`FusionCatcher` uses and ensemble approach consisting of four different methods and four different aligners for identifying the fusion junctions. Each method corresponds to one aligner and the aligners are Bowtie[8], BLAT[9], STAR[16], and Bowtie2[17].

For the first method, which is using information regarding the exon/intron positions (i.e. genome annotation), a database of exon-exon junctions is built, and it contains all the exon-exon junctions for all possible exon-exon combinations for each candidate fusion gene. The unmapped reads are aligned on the database using Bowtie aligner. An unmapped read is counted as evidence for supporting a candidate fusion gene if it is found to (i) map on a exon-exon junction belonging to a candidate fusion gene (i.e. reads R3/2 and R4/2 in Figure 1), and (ii) have its corresponding paired-read mapping on one of the genes forming this candidate fusion gene (i.e. reads R3/1 and R4/1 in Figure 1). The unmapped reads which have the same mapping position on the same exon-exon junction are counted only once. Therefore, the candidate fusion genes which have been found to have counts of such unmapped reads larger than a given threshold will make it to the final list of fusion genes. The role of the first method is to reduce the number of unmapped reads given as input to the next three methods which are more computationally demanding. Also the unmapped reads found to map here are removed from further analysis.

**Figure 1:**
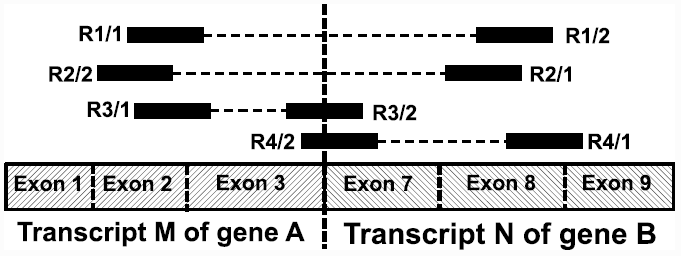
Bowtie’s mappings of reads and their corresponding paired-reads which support the fusion between genes A and B with fusion junction between exon 3 of gene A and exon 7 of gene B. The pairs of reads (R1/1, R1/2) and (R2/1, R2/2) support the fusion and the reads R3/2 and R4/2 support the exon-exon junction of the fusion.

For the next three methods, which are not making use of exon/intron positions (only information regarding start and end positions of genes is used), another database of gene-gene sequences is built (e.g. gene A – gene B shown in Figure 2). The database contains all the gene-gene sequences for each candidate fusion gene. The unmapped reads, which still remain unmapped after aligning them on the exon-exon junctions database using the first method, together with the reads, which are supporting the candidate fusion genes, are aligned further on this database using BLAT aligner[9], STAR aligner[16], and Bowtie2 aligner[17]. A read is counted as evidence for supporting the candidate fusion gene if it is found to (i) map on a gene-gene sequence belonging to a candidate fusion gene such that first part of the read maps on first gene and the second part of the read maps on the second gene (i.e. reads R3/2 and R4/2 in Figure 2), and (ii) have its corresponding paired-read mapping in the transcriptome on one of the genes forming this candidate fusion gene (i.e. reads R4/2 and R5/2 in Figure 2). The reads which have the same mapping positions on the same gene-gene sequence are counted only once, because different reads which share the same start and end position are most likely artefact of PCR process used during sample preparation and sequencing. Therefore, the candidate fusion genes which have been found to have counts of such reads over a given threshold will make it to the final list of fusion genes. The final list of candidate fusion genes encompasses the lists of candidate fusion genes found using all four methods.

**Figure 2:**
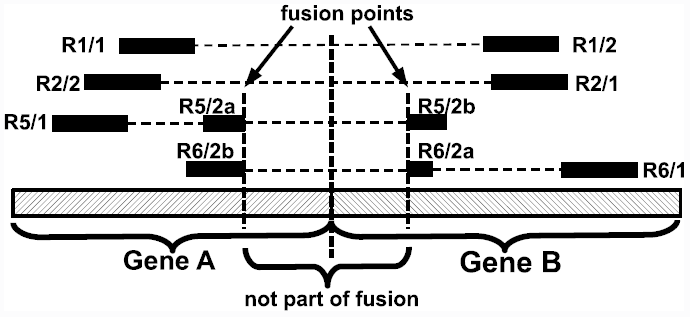
Mappings of reads and their corresponding paired-reads which support the fusion between genes A and B. The pairs of reads (R1/1, R1/2) and (R2/1, R2/2) support the fusion and the reads R5/2 and R6/2 support the junction of the fusion. BLAT, STAR, and Bowtie2 aligners split the read R5/2 into two parts R5/2a and R5/2b and the read R6/2 into R6/2a and R6/2b. The reads R1/1, R1/2, R2/1, R2/2, R5/1, and R6/1 are mapped using Bowtie aligner on transcriptome and the reads R5/2 and R6/2 are mapped using BLAT, STAR, and Bowtie2 aligners on gene sequences.

`FusionCatcher` is written in Python, runs on Linux, and it is freely available under the GPL version 3 license at http://code.google.com/p/fusioncatcher/.

## 3 Results and Discussion

The tumor cell lines SNU-16, KATOIII, and NCI-H716 are known to harbor FGFR2 amplifications. As fusion genes are known to reside on re-arranged areas on the genome, this indicates that the FGFR2 gene is probably a fusion gene partner [11, 14] in these cell lines. This hypothesis is confirmed *in silico* by `FusionCatcher` which found several novel FGFR2 fusions:

- SNU-16 cell line: FGFR2-CD44, FGFR2-PPAPDC1A, FGFR2-MYC, and FGFR2-PDHX [3],
- KATOIII cell line: FGFR2-CEACAM5, FGFR2-ULK4, GCNT3-FGFR2, SNX19-FGFR2, CTNNB1-FGFR2[4], and
- NCI-H716 cell line: COL14A1-FGFR2, FGFR2-COL14A1, PVT1-FGFR2[5].

Additional novel fusion genes have been detected by `FusionCatcher` in HeLa and U87MG cell lines [6, 7].

The breast cancer RNA-sequencing dataset from [2] contains 40 real-time PCR validated fusion genes, of which 27 are published in [2] and 13 in [1]. All 40 fusion genes were detected by `FusionCatcher` and 25 (marked with †in Table 1) were found for the first time by `FusionCatcher` in[1, 2]. In Table 1 is presented a comparison of several fusion genes finders like SOAPFuse[18], ChimeraScan[20], deFuse[27], FusionHunter[28], SnowShoes-FTD[26], and TopHat-Fusion[26] using the same dataset[18]. Also `FusionCatcher` is run (see Table 1) on the same data set using two different Ensembl genome annotations, which are release 61 (based on GRCh38/hg38 assembly) and release 77 (based on GRCh3/hg19 assembly). As expected the genome annotations and genome assemblies used, affect the peformance of fusion genes finding. As shown in Table 1, `FusionCatcher` has the best precision, i.e. *T P/*(*T P* + *FP*), and also competitive sensitivity irrespective to the genome annotation and assembly used.

**Table 1:**
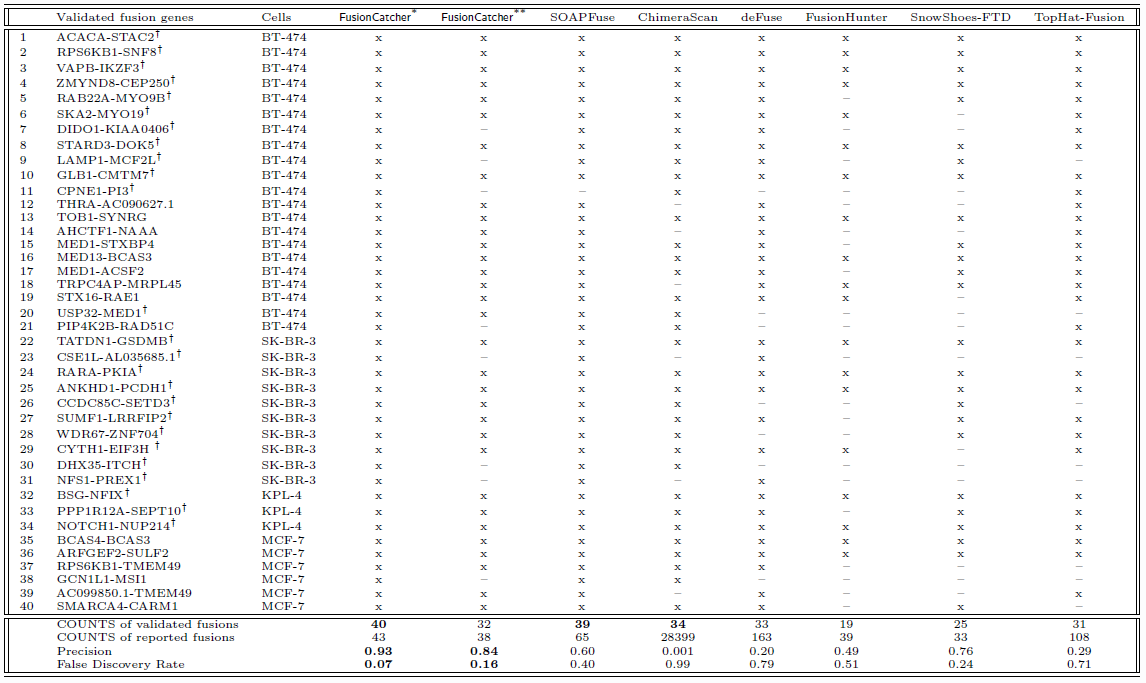
Comparisons of RNA-seq fusion finders on breast cancer cells dataset[2]. `FusionCatcher`* is version 0.91 (i.e. Ensembl release 61 and GRCh37/hg19) from [1, 2]. `FusionCatcher`** is version 0.99.3e (i.e. Ensembl release 77 and GRCh38/hg38). Data regarding SOAPFuse, ChimeraScan, deFuse, FusionHunter, SnowShoes-FTD, and TopHat-Fusion (GRCh37/hg19) are from[18]. Fusion genes marked with ^†^ have been found for the first time by `FusionCatcher`*[1, 2]

In Table 2^3^ is shown another comparison of fusion genes finders on a synthetic spike-in fusion genes dataset from [19], where `FusionCatcher` detects all the synthetic fusion genes.

**Table 2:** Comparisons of fusion genes finders on synthetic spike-in fusion genes dataset (concentration [log10 (pMol)]= -8,57; minimum fusion-supporting read cut off = 2)[19]. `FusionCatcher`** is version 0.99.3e (i.e. Ensembl release 77 and GRCh38/hg38). Data regarding TopHat-Fusion, ChimeraScan, and SnowShoes-FTD (GRCh37/hg19) are from[19].

|  | Spike-in fusion gene | Replicate | FusionCatcher** | TopHat-Fusion | ChimeraScan | SnowShoes-FTD |
| --- | --- | --- | --- | --- | --- | --- |
| 1 | EWSR1-ATF1 | R1 | x | x | x | x |
| 2 | EWSR1-ATF1 | R2 | x | — | x | x |
| 3 | TMPRSS2-ETV1 | R1 | x | — | x | — |
| 4 | TMPRSS2-ETV1 | R2 | x | — | x | — |
| 5 | EWSR1-FLI1 | R1 | x | x | x | x |
| 6 | EWSR1-FLI1 | R2 | x | — | x | x |
| 7 | NTRK3-ETV6 | R1 | x | x | x | x |
| 8 | NTRK3-ETV6 | R2 | x | x | x | x |
| 9 | CD74-ROS1 | R1 | x | x | x | x |
| 10 | CD74-ROS1 | R2 | x | x | x | x |
| 11 | HOOK3-RET | R1 | x | x | x | x |
| 12 | HOOK3-RET | R2 | x | x | x | x |
| 13 | EML4-ALK | R1 | x | — | x | — |
| 14 | EML4-ALK | R2 | x | x | x | — |
| 15 | AKAP9-BRAF | R1 | x | x | x | x |
| 16 | AKAP9-BRAF | R2 | x | x | x | x |
| 17 | BRD4-NUT | R1 | x | x | x | x |
| 18 | BRD4-NUT | R2 | x | — | x | x |
| COUNTS of spike-in fusions |  |  | **18** | 12 | **18** | 14 |

In summary, `FusionCatcher` has the best precision, which translates into in real-time PCR validation rates, for finding fusion genes in the breast cancer RNA-sequencing data set[1, 2, 18] and very good sensitivity in synthetic spike-in fusion genes dataset[19].

## Footnotes

1 November 2014

2 August 2014

3 only samples having concentration of -8,57 are used here because all the other samples are missing the replicates data (November 2014)

